# Fisheries trade and blue nutrient flows in Pacific Island Countries

**DOI:** 10.1101/2025.09.18.677176

**Authors:** Keiko J. Nomura, Steven Mana’oakamai Johnson, Jessica Gephart, Jacob G. Eurich

**Affiliations:** Cooperative Institute for Research in Environmental Sciences, University of Colorado Boulder, Boulder, Colorado, USA; Environmental Data Science Innovation and Impact Lab, University of Colorado Boulder, Boulder, Colorado, USA; Department of Natural Resources and the Environment, Cornell University, Ithaca, New York, USA; School of Aquatic and Fishery Sciences, University of Washington, Seattle, Washington, USA; Environmental Defense Fund, Santa Barbara, California, USA; Marine Science Institute, University of California, Santa Barbara, Santa Barbara, California, USA

**Keywords:** aquatic foods, fisheries governance, food systems, international trade, network analysis, nutritional security

## Abstract

Pacific Island Countries (PICs) are located in highly productive fishing regions that supply nutrient-rich fish to global markets. Marine fisheries are a critical source of protein and essential micronutrients for billions of people worldwide, supporting both local diets and global food security. International trade shapes modern blue food systems, influencing broader distributions and availability of these “blue nutrients.” Yet, the structure of these trade networks, the nutritional composition of exported fisheries, and the implications for local food security remain poorly understood. Using global marine fisheries trade data from 1996-2020 combined with species-level nutrient compositions, we analyzed production, consumption, and nutrient balances for 12 PICs and used network analysis to characterize the structure of international blue nutrient flows. Here we show that many PICs experience persistent net losses of essential nutrients, particularly vitamin B12, protein, and fatty acids. This results in nutrient yields far below population needs, with only a few PICs (Vanuatu and Kiribati) meeting average requirements. Compared to global trade networks, the regional PICs trade is more modular, less reciprocal, and less clustered, implying reliance on a few intermediaries (Papua New Guinea and Fiji). Despite high domestic production and consumption, 54% of blue nutrients are exported from PICs rather than retained locally, leaving local nutritional deficits unaddressed even as PICs supply global seafood markets. Combining trade, nutrient, and network analyses can help inform strategies to increase nutrient retention, strengthen food security, and support resilience for PICs in the modern blue food economy.

## Introduction

Seafood supply chains distribute essential dietary nutrients globally via aquatic food trade (Gephart & Pace, 2015). Aquatic foods (or “blue foods”) encompass marine or freshwater organisms like finfishes, crustaceans, cephalopods, mollusks, seaweeds and algae, and more. They are often critical sources of nutrients such as vitamin B12, iron, and fatty acids which help reduce micronutrient deficiencies and non-communicable diseases, support cardiovascular and brain health, and displace less healthy meats in diets (Golden et al., 2021). As such, there have been many recent calls for better integrating blue foods into global food system solutions and the overall food transformation agenda (Crona et al., 2023; Hicks et al., 2019; Tigchelaar et al., 2022). Aquatic food products consistently rank as one of the most traded food commodities in the world, with global trade volumes equivalent to approximately 38% of production in 2022 (FAO, 2024). Yet, seafood and their nutrients are distributed unevenly via international trade. Lower-income countries tend to export high-value seafood to higher-income countries, while importing lower-value fish for domestic consumption (Lindsay, 2022). Around 36% of nations lose nutrients via fisheries trade, with over 50% of those being Small Island Development States and African nations (Nash et al., 2022). At the same time, undernutrition, micronutrient deficiencies, and overnutrition remain a major challenge in low-income countries (Prentice, 2023) despite rising global fish production per capita (FAO, 2024). Thus, analyzing how trade redistributes fisheries and their associated nutrients – which we call “blue nutrients” – is essential for evaluating modern food systems.

Aquatic foods contribute substantially to the diets of people in Pacific Island Countries (PICs) (Bell et al., 2019; O’Meara et al., 2023), with demand expected to increase through 2050 in many PICs (Teneva et al., 2023). While subsistence fishing and market access support much of the local fish consumption in PICs (Seto et al., 2024), PICs are also among the top exporters of aquatic animal products relative to total exports (FAO, 2024). As a result, a considerable share of blue nutrients is redistributed internationally rather than retained for local use. For instance, much of world’s tuna supply is harvested from the high seas or Exclusive Economic Zones (EEZs) surrounding PICs (Drakou et al., 2018), yet trade and foreign fishing often directs fisheries-derived nutrients toward markets in wealthier, more nutrient-secure countries (Nash et al., 2022). Simultaneously, the region’s caloric needs are highly dependent on imports (Nash et al., 2022), a growing percentage of which are composed of unhealthy food products (Brewer et al., 2023), and diet-related health ailments remain a challenge (Farmery et al., 2021). As smaller, geographically isolated countries with long histories of resource extraction and increasingly liberalized trade policies (Andrew et al., 2022; Campbell, 2015; Havice & Reed, 2012), PICs may exhibit trade patterns that differ from global norms. Understanding these unique trade dynamics is essential for identifying effective interventions to enhance nutrition security and food system resilience in PICs.

In this study, we examined how international aquatic food trade involving PICs influences the distribution and availability of seafood-derived nutrients. We integrated fisheries trade and nutrition data to construct nutrient flow networks at both regional (PICs) and global scales. Specifically, we (1) evaluated how often each country occupied source, exporter, and consumer roles, (2) assessed the proportion of production that is domestically retained versus exported, and (3) calculated net nutrient flows. We then (4) expressed flows as nutrient yields (per capita measures of nutrient availability) to better capture individual-level health implications. Finally, (5) we applied a network analysis to compare the structural features of regional and global trade and assess the positions that PICs occupy in the trade system. Examining these mechanisms allowed us to understand how blue nutrients are traded and the distinct patterns shaping PICs participation in this global system. We conclude by discussing how this integrated perspective on trade, nutrition, and networks can inform policies aimed at improving nutrient retention, trade equity, and food system resilience in the region.

## Methods

### Fisheries Trade and Nutrition Datasets

The Aquatic Resource Trade in Species (ARTIS) consumption database (Gephart et al., 2024) describes international flows of traded aquatic species between the source, exporter, and consumer countries from 1996-2020. Trade is reported by weight (tonnes) primarily at the species-level, and sometimes up to the class-level because ARTIS disaggregates to the lowest scientific name possible based on reported production data and the taxonomic resolution of the codes. The ARTIS database already excludes mammals and seaweeds, and we further focus on marine capture fisheries production for human consumption (rather than species used for fishmeal production or ornamental trade), excluding aquaculture and inland fisheries; our data subset, then, primarily consists of finfish and invertebrates (Gephart et al., 2024). Trade involves transactions where the exporter and consumer differ, while domestic consumption is represented in the database as instances where aquatic species are caught and consumed in the same country.

The ARTIS database was merged with the Aquatic Food Composition Database (AFCD) (Golden, et al., 2021) to assess nutrient flows from international fisheries trade, specifically calcium, fatty acids, iron, protein, vitamin A, vitamin B12, and zinc. The AFCD reports nutrient quantities per unit weight of species. To join ARTIS and AFCD, species-specific nutrient averages were applied to species appearing in both data sets. For higher taxa names appearing in ARTIS, an average for the species falling within that taxa group was applied. For species appearing in ARTIS, but not AFCD, the lowest available taxa average was applied. The resulting “nutrient network” captures the nutrient content of marine fisheries trade flows, while the original trade network represents live species weight. As is standard with apparent consumption data, the ARTIS consumption dataset reports trade in live weight equivalent instead of final food products (e.g., fillet, canned), as the ultimate product form consumed remains unknown in national-level apparent consumption calculations. The datasets used here enable us to convert whole biomass into edible portions, so nutrient flows represent the *total potential nutrients* available for consumption from edible portions, rather than from specific processed products.

Finally, certain fishing arrangements and reporting practices can bias apparent consumption calculations. International fisheries arrangements in the region enable PICs to earn revenue from foreign fishing (mostly for tuna) within their EEZs. Foreign fishing is generally reported in production data under the flag of the vessel and may not fall within a country’s trade data reporting. Consequently, these flows of fishery products out of the country are often not captured in trade data. Meanwhile, joint ventures result in production data attributed to the host nation, but may still not be captured in export data, resulting in artificially high apparent consumption calculations. Additionally, port stops by foreign vessels may be inconsistently reported as re-exports or not reported at all as products in transit through a country are optional to report as trade. The data also only includes countries, with territories aggregated into the sovereign state due to data availability considerations such as inconsistent territorial reporting. Lastly, small-scale fisheries (SSF) are often underreported in production data, which can underestimate domestic production estimates, especially species primarily targeted by SSF for local consumption. Despite these considerations, the constructed trade networks are still useful, and our interpretation accounts for these limitations.

### Species Composition, Nutrient Flows, and Trade Roles

Here, we examine twelve sovereign PICs: Fiji (FJI), Federated States of Micronesia (FSM), Kiribati (KIR), Marshall Islands (MHL), Nauru (NRU), Palau (PLW), Papua New Guinea (PNG), Solomon Islands (SLB), Tonga (TON), Tuvalu (TUV), Vanuatu (VUT), and Samoa (WSM). Pacific Island Territories (e.g., Cook Islands, Niue, French Polynesia) are not analyzed here due to the aggregation challenges discussed above. To compare global versus regional trade dynamics involving PICs, we sampled a subset of the trade network by gathering all transactions that involve a PIC either as source, exporter, or consumer. For example, a trade pathway where Fiji is the source, China the exporter, and USA the consumer would be included in the PICs network.

The resulting regional PICs data was used to examine properties of the nutrient networks, like prominently traded nutrients and species, which roles countries occupy (i.e., source, exporter, consumer), national net changes in nutrient yields, and to what extent domestic production contributes to within-country nutrient retention. For each PIC, we calculated *consumption origin ratios* that quantify the proportion of consumed nutrients that are domestically produced versus imported. We also calculated *production fate ratios* that quantify the proportion of nutrients from PICs’ fisheries production that are consumed domestically versus exported for foreign consumption.

### Trade Balances and Population-Level Nutrient Yields

The absolute quantities of different nutrients traded between countries are difficult to compare directly because they are measured in incompatible units (i.e., grams, micrograms, milligrams). To address this, we calculate a *richness ratio* for each nutrient flow by dividing nutrient quantity by its Recommended Nutrient Intake (RNI), expressing nutrient flows as daily dietary fulfillments (similar to Golden et al. (2021)). For example, a protein richness ratio of 25 flowing from country A to country B indicates that 25 daily dietary protein recommendations are contained in the marine fisheries traded from A to B. For these calculations, RNI values for reproductive aged women (19-50 years old), a population often vulnerable to undernutrition, were sourced from World Bank (National Institutes of Health Office of Dietary Supplements, 2024). For each trade flow, we also calculated an average richness ratio across all individual nutrients, following Golden et al. (2021). Positive richness ratios indicate positive trade balances, where countries have a net-inflow of nutrients; negative ratio values indicate a negative trade balance, or net out-flow via trade.

Because a country’s nutrient requirements depend on its population size, we calculated *nutrient yields* for each country by dividing richness ratios by national population and time, following Nash et al. (2022). Nutrient yields are expressed in units of RNIs per capita per day, where a value of 1 indicates that approximately every person receives an adequate supply of that nutrient. Annual population data for most countries were obtained from the World Bank for 1996 to 2020 (World Bank, 2024). For Pacific Island Countries, more accurate population estimates were sourced from the Pacific Data Hub (Pacific Community, 2024b), while Taiwan’s population data were collected from its National Statistics database (Taiwan Department of Statistics, 2024), as it was missing from the World Bank dataset. Population data were used to calculate annual and time-averaged nutrient yields for each country, as well as aggregate yields across nutrients, enabling assessment of net surpluses, deficits, and population-level impacts. All analyses were performed in R (R Core Team, 2024) using the *rsdmx* (Blondel, 2025) and *WDI* (Arel-Bundock, 2025) for population data access, and *tidyverse* (Wickham et al., 2019) for data manipulation.

### Trade Network Analysis

Nutrient flow networks were constructed at both regional (PICs) and global scales. In both networks, nodes represent countries; weighted, directed edges represent nutrient flows that signify the quantity (in RNIs) and direction of trade. Edges were collapsed into bilateral country-to-country nutrient flows (e.g., source-to-exporter and exporter-to-consumer) rather than modeled as full tripartite pathways (e.g., source-exporter-consumer). Each edge thus represents the cumulative nutrient flow between two countries, regardless of role. Consistent with prior regional sampling, the regional PICs network consists of all trade transactions involving a PIC in any role. By contrast, the global network encompasses the full global ARTIS dataset, including trade pathways with non-PIC actors. This allows us to compare the structure of nutrient flows in PICs with global patterns.

To characterize the structure of nutrient flows, we compared network-wide and node-specific metrics between regional and global scales. Networks have been widely used to analyze structural patterns in economic systems such as trade (Fagiolo et al., 2010; Serrano et al., 2007), including seafood trade (Gephart et al., 2016; Gephart & Pace, 2015). This approach enables evaluation of both the entire trade structure through network-wide metrics and country-level positions through node-specific metrics. In the context of nutrient flows, network analysis can illuminate blue food security by quantifying distribution efficiency, national in-and out-flows of nutrients, and the centrality of countries within the system. Specifically, we analyzed a suite of network-wide metrics: edge density, average path length, assortativity, centralization, diameter, modularity, reciprocity, and transitivity (Barrat et al., 2004; Fagiolo, 2007; Newman, 2010). At the node-level, we calculated each node’s degree, strength, betweenness and closeness centrality, and local clustering coefficients (Barrat et al., 2004; Freeman, 1977; Opsahl et al., 2010) **(Figure 1**). See Figure 1 for explanations of each metric and its interpretation in a trade context. By comparing these metrics across scales, we aim to identify structural patterns specific to the regional network in contrast to the global system.

**Figure 1.**
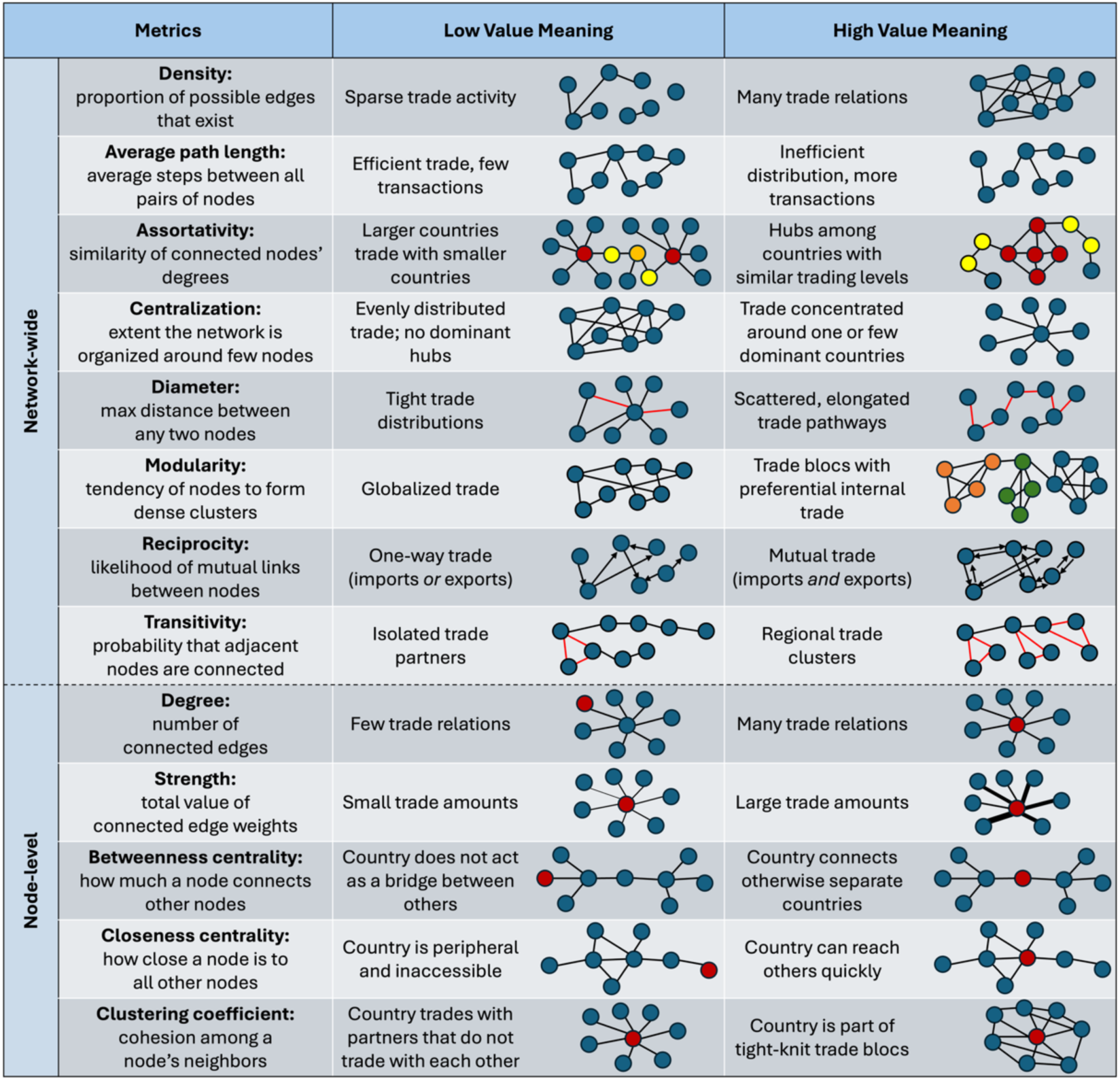
Network metrics used to characterize blue nutrient trade networks. The left column defines each metric. The middle and right columns interpret and illustrate low and high values, respectively, in a trade context. Network-wide metrics describe the overall structure of the network, while node-level metrics characterize individual countries’ positions within it. Red nodes indicate the focal country in node-level diagrams.

To assess whether the empirical nutrient networks’ features were statistically distinct from random expectations, we employed two complementary null model strategies. First, we directly compared the empirical PICs and global nutrient networks by generating 100 randomized replicates of each using a directed degree-preserving configuration model (Fosdick et al., 2018). This ensured that each replicated network maintained the same network-wide degree patterns as the empirical network, constraining the possible values of network metrics. We then calculated most of the same network-wide and node-level metrics described above (excluding degree and centralization, which are fixed by the imposed degree sequence and therefore not suitable for statistical comparison). Then, we performed two-sample t-tests on the regional and global simulated metrics to evaluate any significant structural differences.

Second, we similarly examined whether the structure of the PICs network alone is significant. Using a Markov Chain Monte Carlo edge-swapping algorithm (Fosdick et al., 2018), we sampled uniformly from the space of networks with the same degree sequence. We calculated the same network-wide and node-level metrics for each replicate, then used z-scores and two-tailed p-values to assess whether the empirical values significantly deviated from the null distributions. All network analyses, statistical comparisons, and visualizations were performed in R (R Core Team, 2024) using the *igraph* (Csárdi et al., 2025) and *ggraph (Pedersen, 2024)* packages.

## Results

### Species Composition, Nutrient Flows, and Trade Roles

Between 1996 and 2020, the PICs fisheries trade network involved approximately 16.9 million tonnes across 1,603 species or groups from marine capture fisheries. Four large pelagic species totaled nearly 71% of the overall tonnage and 65% of the transactions traded (**Figures S1 & S2**): skipjack tuna (*Katsuwonus pelamis*, ∼7.7 million tonnes; 122,000 transactions), yellowfin tuna (*Thunnus albacares*, ∼3.0 million tonnes; 146,000 transactions), bigeye tuna (*Thunnus obesus,* ∼679,000 tonnes; 111,000 transactions), and albacore tuna (*Thunnus alalunga*, ∼600,000 tonnes; 65,000 transactions). The broader taxonomic group *Actinopterygii* spp. contributed another 8.7% of total trade weight, while a few small to medium pelagics (*Trachurus murphyi*, *Scomber japonicus*, *Sardinops sagax*) contributed 9.4%. Trade frequency and weight were highly correlated (Pearson r = 0.70, Spearman ρ = 0.98, both p < 0.001), indicating dominance of high-volume, high-frequency trade of a small number of key species.

Trade composition varied by role. The top species for source and exporter roles were 52% similar, suggesting some alignment between harvest and export. However, approximately 84% of top source species appeared in the consumer rankings, suggesting that countries often consume the fish that they harvest. Among the top 25 species, overlap ranged from 52-84%, indicating both a core group of commonly traded species (e.g., skipjack tuna) and role-specific specializations.

Nutrient flows (in RNIs) were dominated by vitamin B12 (145.2 billion, 70.5%), fatty acids (26.6 billion, 13.4%), and protein (25.6 billion, 11.2%), followed by iron (4.7 billion, 2.2%), zinc (3.9 billion, 1.8%), calcium (1.5 billion, <1%), and vitamin A (1.1 billion, <1%). These patterns reflect the nutrient profiles of commonly traded species. Large pelagic fish contribute high levels of B12, fatty acids, and protein, while species typically consumed whole or canned with bones and organs (often small pelagics; e.g., *Sardinella*, *Engraulis*, and *Clupea* spp.) contribute more calcium (Isaacs, 2016). Nutrient flows were strongly correlated with trade weight (r² > 0.89 for most nutrients, except calcium at r² = 0.73; all p < 0.001), reflecting the nutrient profiles of traded species.

Country roles differed regionally and globally. In the regional network, 9 PICs ranked among the top source countries **(Figure S3).** Large non-PIC sources included China, Indonesia, New Zealand, and the United States, illustrating both major fishing countries and the geographic bias of nearby trading partners. Fiji and Papua New Guinea are prominently ranked as regional exporters. The top ten largest consumers are all PICs, with Fiji, Papua New Guinea, and Samoa occupying the top three slots. The largest non-PIC exporters in the regional trade network are China, Thailand, the United States, and Taiwan, and the largest non-PIC consumers are Japan, the United States, China, and Australia. Of the total RNIs flowing in the regional trade system, domestic consumption and production within PICs comprise ∼35%. Major trade pathways originated mainly from PICs, followed by Asia (**Figure 2**). The absence of PICs from the top 30 for any role in global trade illustrates their small global footprint.

**Figure 2.**
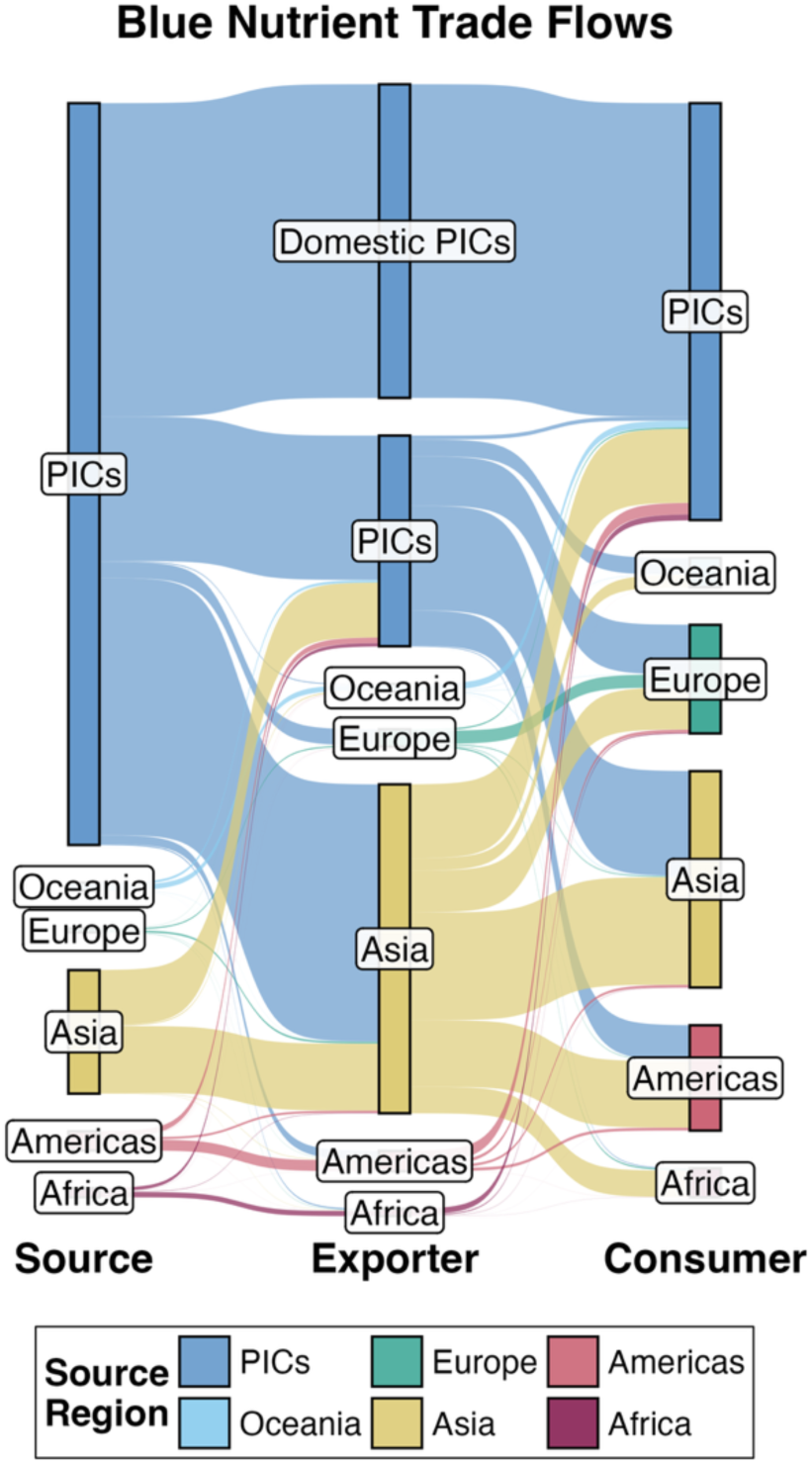
Sankey diagram of fisheries nutrient trade flows (in RNIs) by region from Source (left) to Exporter (middle) to Consumer (right). Flows are shown at the continent level, with PICs as a separate category to highlight their trade links. “Domestic PICs” in the Exporter column denotes production and consumption within the same PIC. Block height reflects each region’s proportional contribution, flow color indicates the Source, and flow thickness represents ratios in RNIs.

### Nutrient Trade Balances and Population-Level Yields

Net flows reflect gains through domestic production and imports, minus losses from exports. For most PICs, domestic sources were the primary sources of positive flows (**Figure S4**). Only Fiji, Palau, and Samoa received more fisheries trade through imports than they produced; these three were also the only ones with imports exceeding exports. Half of PICs had net negative outflows and half had positive inflows, ranging from – 878,000 tonnes (Marshall Islands) to 470,000 tonnes (Kiribati). Among countries with net positive inflows, domestic production accounted for most gains, from 56% for Samoa to domestic production exceeding net flows by over 300% in Vanuatu.

Net trade flows were converted to nutrient richness ratios, showing gains or losses in RNIs from 1996 to 2020. As with trade weight, countries varied in both the direction and magnitude of nutrient flows (**Figure S5**). Fiji, Kiribati, Palau, Samoa, Tonga, and Vanuatu had positive richness ratios across all seven nutrients, while the Marshall Islands, the Federated States of Micronesia, Papua New Guinea, Solomon Islands, and Tuvalu showed losses across all nutrients. The Marshall Islands experienced the largest deficits, ranging from 55.2 million RNIs of calcium lost to 27.1 billion RNIs of vitamin B12 lost. Kiribati, Solomon Islands, and Vanuatu were negative across all nutrients except calcium. All countries’ nutrient flow direction matched their trade weight trends.

While richness ratios capture total gains or losses, nutrient yields translate these flows into per capita dietary contributions. Only two PICs had aggregate nutrient yields greater than 1 (Kiribati and Vanuatu), meaning most countries did not meet daily RNI needs through trade. Five of twelve PICs had negative values (**Figure 3**), ranging from approximately –0.03 in Papua New Guinea to –10.7 in the Marshall Islands, signifying that individuals in these countries *lose* approximately 0.03 to 10.7 aggregate RNIs each day from trade. In contrast, nutrient-positive countries gained between 28% (Palau) and 98% (Samoa) of daily RNIs per person. Vitamin B12 flows had by far the largest impact of any of the individual nutrients. Overall, most countries did not meet adequate nutrient intake through trade, with many experiencing negative yields.

**Figure 3.**
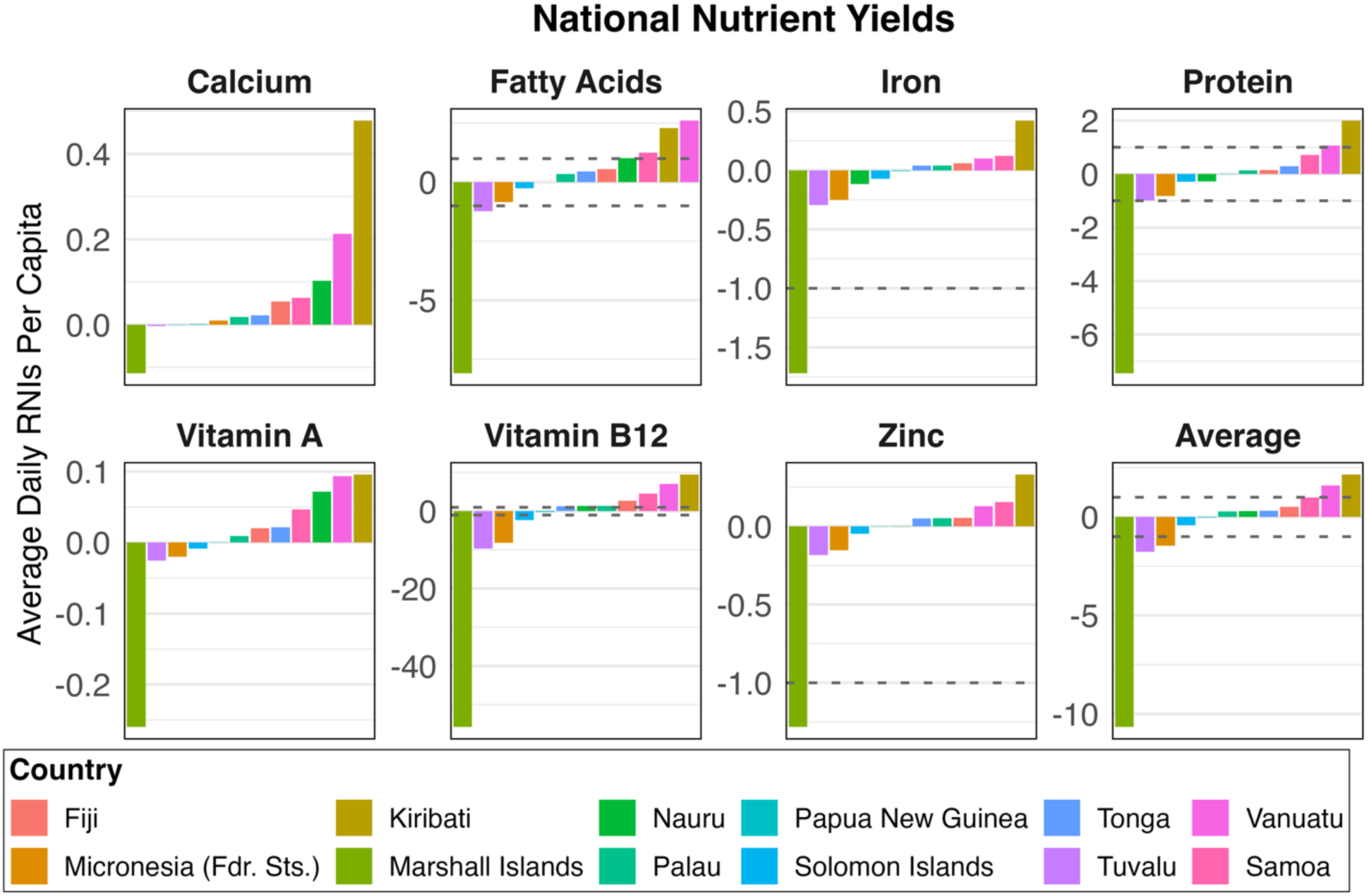
Per capita nutrient yields from fisheries trade in PICs, calculated annually as nutrient RNIs per person per 365 days, then averaged over the time period. Each panel depicts a PIC, with colored bars showing individual RNIs or average nutrient yield quantities. Positive values indicate net nutrient gains, and negative values indicate net losses. The dotted line at y = 1 marks that a country’s entire population’s RNI needs are met through trade, values below 1 indicate partial or no fulfillment, and values below -1 reflect losses greater than daily needs. For example, y = -2 means the nutrients lost are equivalent to double the population’s daily needs.

### Domestic Production and Consumption

Domestic production and consumption play a key role in PICs nutrient retention. On average, ∼74% of RNIs consumed in the region are produced domestically rather than imported (**Figure 4**). Fiji had the lowest domestic consumption origin ratio at around 33%, while Kiribati had the highest at about 98%. On average, only ∼46% of domestically produced seafood is consumed locally, with ∼54% exported. Tonga retained the most domestically produced fish, with a domestic production fate ratio of about 79%. The Marshall Islands had the highest exported production ratio at roughly 85%, despite also having a high domestic consumption origin ratio. Paired with high per capita seafood consumption in PICs (Sharp & Andrew, 2024), these ratios suggest that most PICs are highly reliant on domestic fisheries for their own consumption, yet prioritization of fish for export markets also plays a role (Campbell, 2015). As a result, they simultaneously export over half of their catch and rely heavily on domestic production to meet local demand.

**Figure 4.**
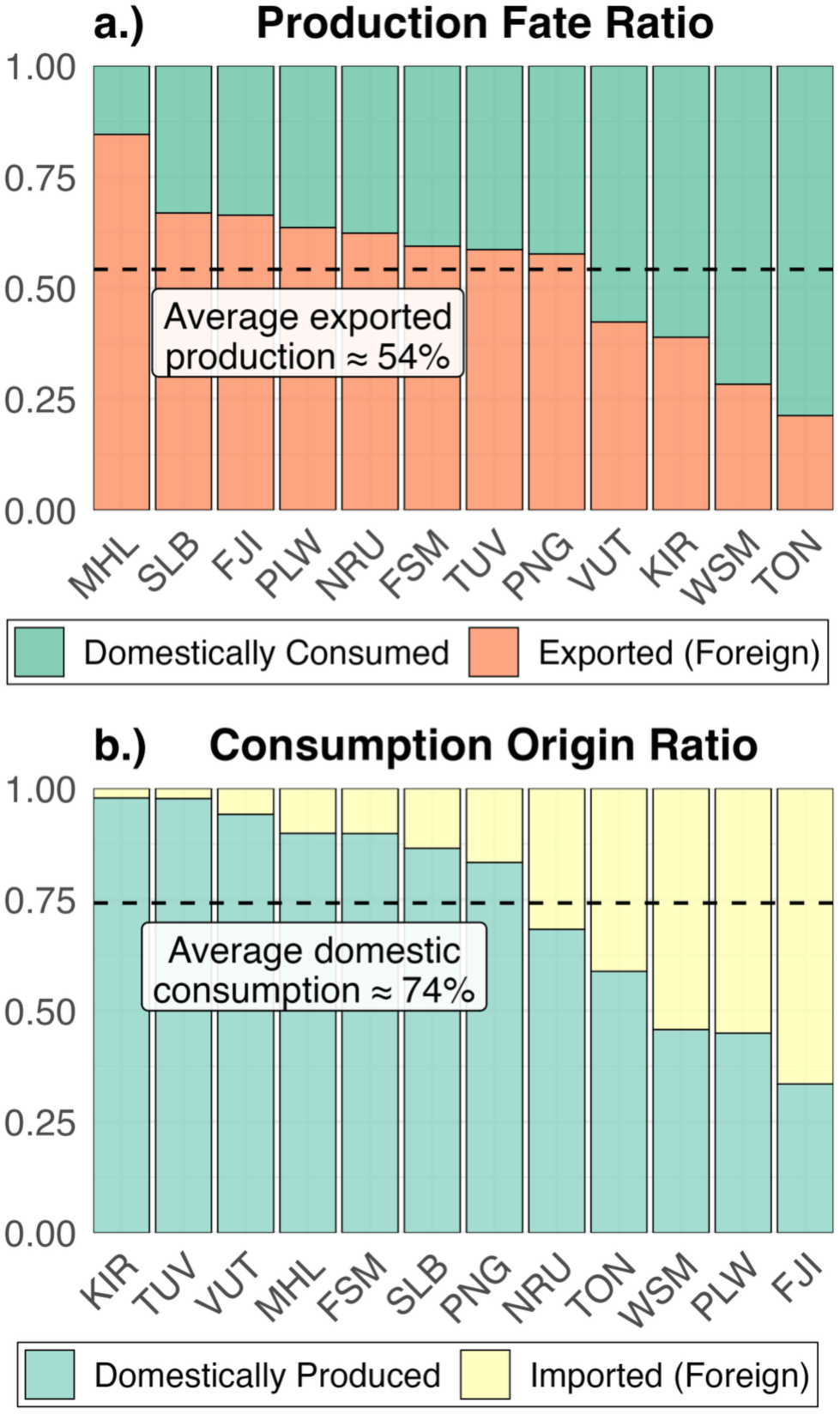
Production fate ratios (a) and consumption origin ratios (b) for all twelve PICs. Bars show each country (x-axis) and proportion (y-axis) in species tonnes. In a.), teal indicates the proportion of catch domestically consumed, orange indicates the share exported, and the dotted line marks the average exported (∼54%). In b.), blue shows the proportion of consumed seafood that is domestically produced, yellow shows the share imported, and the dotted line marks the average consumption domestically produced (∼74%). PICs are abbreviated as follows: Fiji (FJI), Federated States of Micronesia (FSM), Kiribati (KIR), Marshall Islands (MHL), Nauru (NRU), Palau (PLW), Papua New Guinea (PNG), Solomon Islands (SLB), Tonga (TON), Tuvalu (TUV), Vanuatu (VUT), and Samoa (WSM).

### Global and Regional Trade Network Comparisons

Network metrics differed substantially between the regional (PICs) and global seafood trade networks, though both included 192 countries (**Table S1**). The global network had nearly twice as many trade flows (∼15,700 vs. ∼8,900), with higher density (0.43 vs. 0.24) and average degree (163 vs. 92). Both degree distributions were roughly normal, though the PICs network was more right-skewed, indicating fewer dominant countries and a generally sparser network (**Figure S6**).

These structural differences influenced how nutrients move, how trade is organized, and how resilient the system may be to disruption. The global network had a longer average path length (40,600 vs. 6,400) and a larger diameter (3.6 million vs. 537,000), implying that global trade is more indirect and spread across longer chains of intermediaries, whereas the PICs network is relatively more compact. Higher centralization in PICs (0.61 vs. 0.53) reflects greater reliance on fewer key hubs, which may increase vulnerability to disruption in key trade routes. While modularity was higher in the regional network (0.25 vs. 0.12), clustering (0.54 vs. 0.71) and reciprocity (0.62 vs. 0.70) were lower. These patterns imply that, although the PICs network has delineated trade communities, it lacks the tightly knit, reciprocal relationships found in the global system, aligning with the region’s overall net outflow of nutrients. Our analysis also confirms a more modular and asymmetric trade structure (**Table S2**). Notably, reciprocity exceeded null expectations, suggesting a greater-than-random tendency for mutual trade ties within the region (**Table S3**). Overall, these findings suggest that non-random factors such as geographic isolation, limited trade partners, and policy arrangements may shape the PICs trade network and lead to structural differences from global patterns.

### Country Positions in the Trade Network

Several PICs emerged as structurally important in the regional trade network (**Figure 5**). Papua New Guinea had the highest betweenness centrality, acting as a key conduit for nutrient flows but with strongly negative net strength, reflecting major outbound flows. This aligns with its status as the largest exporting PIC and the third most frequent source in the region. Fiji ranked the second highest in betweenness and had the highest total degree, reinforcing its role as a central trade hub. Unlike Papua New Guinea and other central nodes with negative outflows (e.g., Kiribati, Nauru), Fiji had a positive net strength, indicating that it functions both as a connector and as a recipient of nutrients. These roles reflect prior findings that identify Fiji and Papua New Guinea as regional tuna processing and re-export centers, supported by infrastructure and ties to major global supply chains (Andrew et al., 2022; Bell et al., 2019; Farmery et al., 2020).

**Figure 5.**
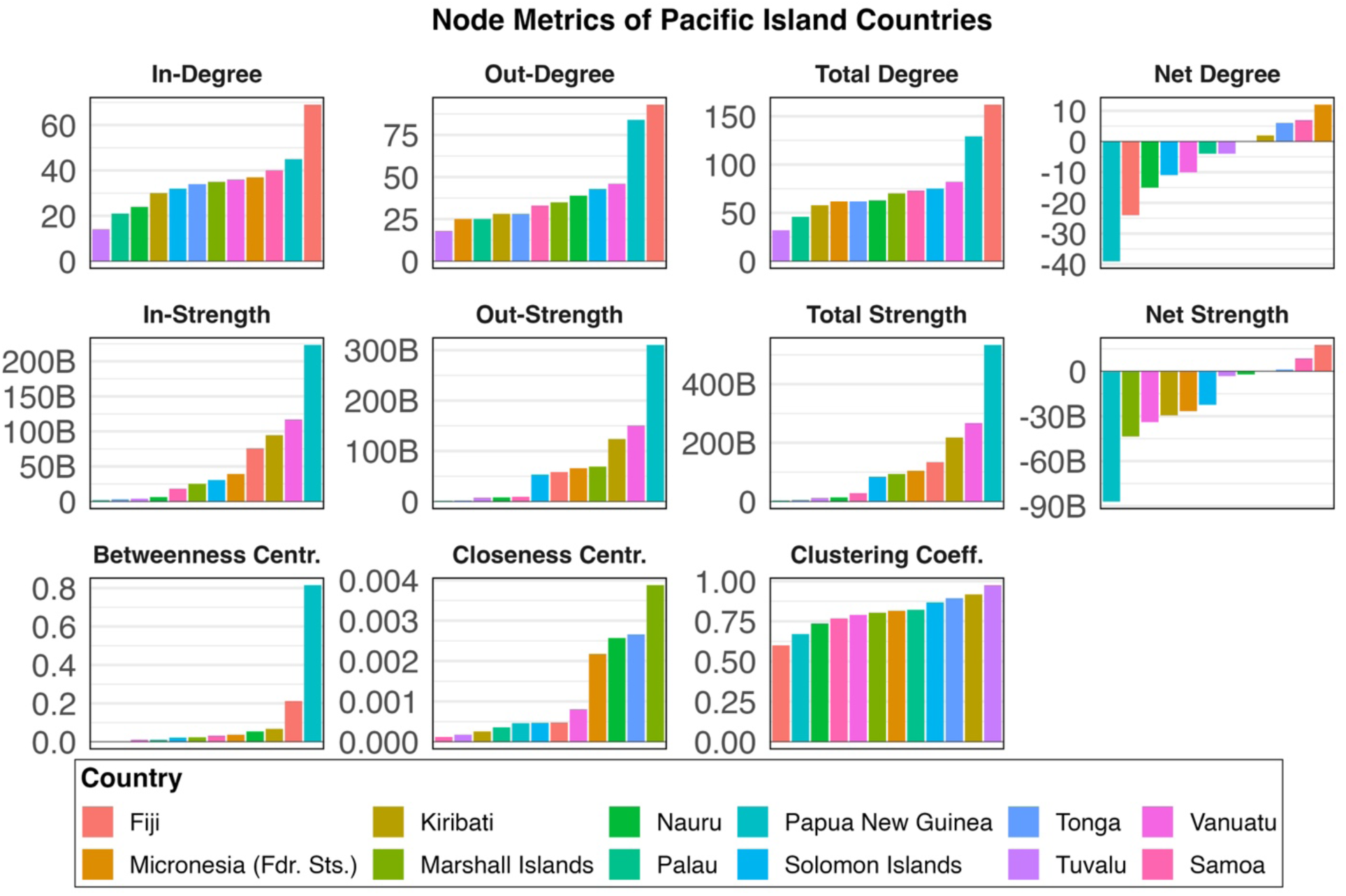
Node-level metrics for all PICs in the regional trade network. Each panel displays a different node-level metric for the PICs, which are represented by different colored bars. The top row displays degree values (i.e., the number of edges), while the second row displays strength values (i.e., the sum of edges). For these metrics, “in” and “out” specifies the directionality of the edges towards or away from the node; total and net values represent the absolute sum and the difference, respectively. The bottom row displays centrality (i.e., the node’s connectedness) and clustering coefficient values (i.e., how clustered the node is). See Table 2 for more detailed explanations of metrics.

Samoa had low betweenness and the lowest closeness, underscoring its peripheral position, yet it retained a positive net balance. Tonga likewise showed zero betweenness but relatively high closeness, suggesting it is accessible without acting as a broker, and it too held a modest surplus. In contrast, the Marshall Islands stood out with the highest closeness among PICs but the most severe negative trade balance, illustrating how structural accessibility can coincide with major nutrient losses. Statistical comparison with null models reinforced these findings: several PICs, including Papua New Guinea, Fiji, Kiribati, and Nauru, had significantly higher betweenness than expected, underscoring their role as regional connectors. Yet, several of these same countries (e.g., Nauru, Samoa, Palau, Micronesia, and the Marshall Islands) showed significantly lower clustering than null expectations, indicating outward-oriented ties with little redundancy. Overall, PICs occupy highly asymmetric roles in the network: some serve as bridges and exporters (e.g., Papua New Guinea, Marshall Islands), others as hubs and consumers (e.g., Fiji), and others remain peripheral but manage modest nutrient surpluses (e.g., Samoa, Tonga).

## Discussion

Seafood trade plays a critical role in distributing blue nutrients globally. However, for PICs, the structure and outcomes of trade raise important questions about local nutrient availability and equity. This paper explores fisheries trade and its implications for nutrient flows and availability in PICs. We find that PICs primarily have net negative fisheries trade balances in both weight and nutrients. In many cases, nutrient losses equal or exceed countries’ daily population requirements. While per capita seafood consumption remains high within PICs, they produce far more than they consume, reinforcing their role as a global supplier of seafood-derived nutrients. PICs are a major source of tuna, with much of this production exported to Asian and European countries. Structurally, fisheries trade involving PICs is comparatively sparse and hierarchical, relying on a limited set of partners rather than densely interconnected trade webs present at the global level. Fiji and Papua New Guinea serve as key distributors, though in contrasting ways: Fiji functions as a central hub with a positive nutrient balance, while Papua New Guinea acts as a major broker but experiences large net losses. By understanding the nutritional imbalances in seafood trade, this work lays a foundation for more just and nutrition-sensitive blue food systems in the Pacific. Below we outline the findings and implications of our analysis of fisheries trade in the PICs, focusing on how current patterns influence nutrient flows, economic benefits, and food system equity.

Tuna dominates fisheries trade from PICs, with most of the global supply being harvested through international and foreign fishing activities in the western central Pacific Ocean (Williams & Ruaia, 2023), and ultimately exported to wealthier and/or more nutrient-secure countries (Nash et al., 2022). Catches flow into several global markets for canned, frozen, or fresh products (Trade Flow Analysis of Pacific Tuna Fisheries, 2023). Despite the economic benefits of tuna fisheries for PICs, redirecting these resources towards local access and consumption could enhance nutritional security (Bell et al., 2015). For instance, shelf-stable canned tuna contributes to increasing proportions of recommended dietary fish intakes for several PICs like Fiji and Solomon Islands, yet better access will be needed to meet population needs across all PICs (Bell et al., 2019). In addition to albacore, yellowfin, and skipjack tuna, mackerel and sardines are also common species used for canning (Bell et al., 2019; Farmery et al., 2020), and all prominently traded in our analysis (∼3.3% traded weight was mackerel; ∼2.9% for sardines).

These dynamics also reflect structural economic inequities in seafood value chains. Despite regional tuna productivity, PICs are increasingly reliant on canned tuna imports primarily from China and Thailand, even as major canneries exist in Fiji, Papua New Guinea, and the Solomon Islands (Farmery et al., 2020). Most processing occurs offshore in Asia or in foreign-owned facilities on PICs. These arrangements result in missed potential revenue from value-added processing and generate fewer opportunities for local workers, and overall limited local economic benefit (Havice & Reed, 2012), consistent with broader corporate concentration of power in fisheries and food systems (Clapp et al., 2025; Folke et al., 2019).

While our analysis focused on traded fishery products, nutrients from aquatic foods can also leave national waters through distant water fishing (DWF) without entering formal trade. Distant water fishing nations (DWFNs) may pay access fees to coastal states where they harvest surplus allowable catch beyond what the State can harvest itself. For instance, Pacific Island nations participate in the Parties to the Nauru Agreement, where DWFNs harvest highly mobile and valuable species like tuna (Palau Arrangement, 2024). Globally, foreign fishing by DWFNs is estimated to relocate about 1.5 times more fisheries-derived nutrients than trade (Nash et al., 2022). PICs are particularly vulnerable to these nutrient flows: some are highly sensitive to blue nutrient losses (Solomon Islands, Micronesia, Tuvalu, Papua New Guinea), others are highly exposed to shifts in trade and foreign fishing (Vanuatu, Samoa, Tonga, Fiji), and Kiribati faces both (Nash et al., 2022). Transparency in DWF and fisheries access arrangements could help clarify how foreign fishing fleets affect blue nutrient distributions, but processes such as transshipment and flags of convenience complicate efforts to accurately trace and quantify these flows. Limited transparency also raises questions about whether resource owners capture the full economic benefits or if these benefits are equitably distributed (Campling et al., 2024; Skerritt, 2024). For example, Chinese and Taiwanese DWF fleets have re-flagged vessels under PICs to exploit discounted license fees and fishing exemptions meant for SIDS’ development (Campling et al., 2024). These dynamics favor DWFNs and hinder the redistribution of benefits to resource-owning states. Thus, when designing policies to ensure nutrition-sensitive management of aquatic resources, flows of traded products must be considered alongside products derived from DWF that do not enter trade.

Given the rising imports of unhealthy foods into PICs (Brewer et al., 2023), there is a clear need to supplement economically-driven trade policies with human health objectives to improve food systems. Globalized trade policies are contributing to a “nutrition transition” toward more Western-style diets characterized by less traditional foods and more caloric, sugary, and processed foods (Golden, et al., 2021; Hawkes, 2006; Popkin & Gordon-Larsen, 2004). In turn, this may be contributing to high rates of obesity and other diet-related non-communicable diseases (Andrew et al., 2022; Sahal Estimé et al., 2014). We found that most PICs experienced substantial blue nutrient losses from fisheries trade, especially in vitamin B12 and, in some cases, calcium. These micronutrients are essential for neurological function and bone health (Phogat et al., 2022), and improved intake could address deficiencies, support healthier aging, and reduce overall disease burden (Golden, et al., 2021). This is especially true given the prominence of health issues and domestic pushes to enhance blue food retention and security.

The different patterns between the global and PICs regional trade networks underscore the need for a more PIC-focused lens on food trade policy. The PICs network is not only smaller, but also distinct in being more modular and less reciprocal, with clearer community divisions and fewer redundant links. As such, certain PICs are reliant on a small number of intermediaries. Fiji and Papua New Guinea, for example, stand out as major exporters with key bridging roles in facilitating flows between countries with otherwise limited direct trade. Their prominence may reflect geographic accessibility, processing infrastructure, and established trade relations (Trade Flow Analysis of Pacific Tuna Fisheries, 2023). Indeed, enhancing intra-regional supply chains could support equitable distribution systems to Pacific Islanders (O’Meara, et al. 2023), and using network-informed interventions can target management (e.g., infrastructure investments or diplomatic facilitation) at critical connectors. Notably, Fiji and Papua New Guinea, the region’s largest economies, declined to sign the Pacific Agreement on Closer Economic Relations Plus in 2017 that would strengthen trade with New Zealand and Australia, citing concerns over policy autonomy and tariff impacts (Morgan, 2018). Their central role in the regional trade network, as shown in our analysis, may have also influenced this decision. In contrast, more peripheral countries like Tuvalu could benefit from greater regional cooperation and improved market access. These examples illustrate how uniform trade policies can overlook the diverse needs of PICs, risking marginalization of smaller states in global food system dialogues.

A broader suite of domestic policies has been proposed across the region to support more resilient, nutrition-sensitive food systems. Many focus on better integration of fisheries and food objectives, such as reinvesting tuna license revenue into supply chain infrastructure (Bell et al., 2019), increasing domestic fishing capacity (Bell et al., 2015; Berry et al., 2022), managing fisheries for maximum nutrient yields (Robinson et al., 2022) or climate resilience (Eurich et al., 2024), and supporting SSF through secure tenure rights and community-based management (Basurto et al., 2025). Since trade and fishing substantially redistribute blue nutrients away from PICs (Nash et al., 2022), these cross-sectoral approaches offer opportunities for broader benefits. Other measures include expanding aquaculture production, nutrition education, and public procurement programs to improve access to blue foods. These initiatives reflect a growing recognition that food security needs coordinated action across all sectors. Scaling these efforts will require stronger governance, financing, and regional collaboration to overcome infrastructural and capacity constraints, especially in more remote or import-dependent countries.

National priorities are reflected, and further differentiated, through Food Systems Pathways (FSPs) developed by the twelve PICs analyzed here (Berry et al., 2022). Most FSPs mention reducing reliance on imported goods, but only half address blue foods and fisheries directly (Berry et al., 2022). Among those that do, the strategies are diverse: Kiribati, for example, is working to improve local fisheries value chains to replace imported goods, while the Marshall Islands is exploring eco-labelling and ocean-to-plate marketing strategies. However, most FSPs still lack plans to address biodiversity, food marketing, climate change, and traditional knowledge, all of which affect blue food outcomes (O’Meara et al., 2023). Despite their social, ecological, and economic significance, fisheries in PICs remain underemphasized in national food security planning.

Although international trade shapes seafood flows at the global scale, it is only one aspect of blue food security. In many PICs, a large proportion of consumed seafood comes from coastal fisheries and smaller-scale operations (Wabnitz et al., 2023). These local seafood resources often flow through non-market acquisition pathways like home production and gifting (Seto et al., 2024). Access to different species varies: wealthier households consume more pelagic species, while low-technology households rely on coastal species like crabs and shellfish (Seto et al., 2024). Another factor shaping local seafood access is tourism, which is a major part of many PIC economies but can also temporarily boost demand and divert fish to restaurants catering to visitors (Colette C.C. Wabnitz, 2019). Proposed strategies, such as reserving reef fish for locals and serving pelagic species to tourists, highlight the need to integrate blue food and broader economic policies to reduce pressure on local supply (Birkeland, 2017). This demonstrates the limits of our trade-based analysis, as the international-level approach cannot capture how traded fish actually reach different segments of the population. Factors like price, infrastructure, domestic markets, preference, demographics, and urban-rural dynamics will influence within-country distributions (Charlton et al., 2016; Wabnitz et al., 2023). Integrating data on local production and consumption (e.g., Household Income and Expenditure Surveys (Pacific Community, 2024a)), could provide a more complete view of blue food dynamics across scales, supporting food system transformations through context-specific understandings of both top-down and bottom-up patterns (Conti et al., 2025).

This analysis demonstrates that international fisheries trade facilitates substantial blue nutrient losses from PICs, potentially reinforcing structural inequities in global seafood systems. Addressing these imbalances will require multi-scale policies that prioritize local value retention, strengthen regional trade, and invest in supply chain infrastructure. Future work should also integrate within-country access, non-market seafood sources, dietary transitions, and the health impacts of rising agricultural and processed food imports. Ultimately, achieving more equitable and resilient blue food systems in the PICs region will depend on coordinated approaches that align international trade with local nutrition and economic priorities, while recognizing the full diversity of food system pathways across the region.

## Acknowledgements

This study was supported by the National Science Foundation award DBI 2153040, Environmental Data Science Innovation and Impact Lab (ESIIL): Accelerating Discovery by Fostering an Open and Diverse Earth Data Revolution.

## Supplementary Information

**Figure S1.**
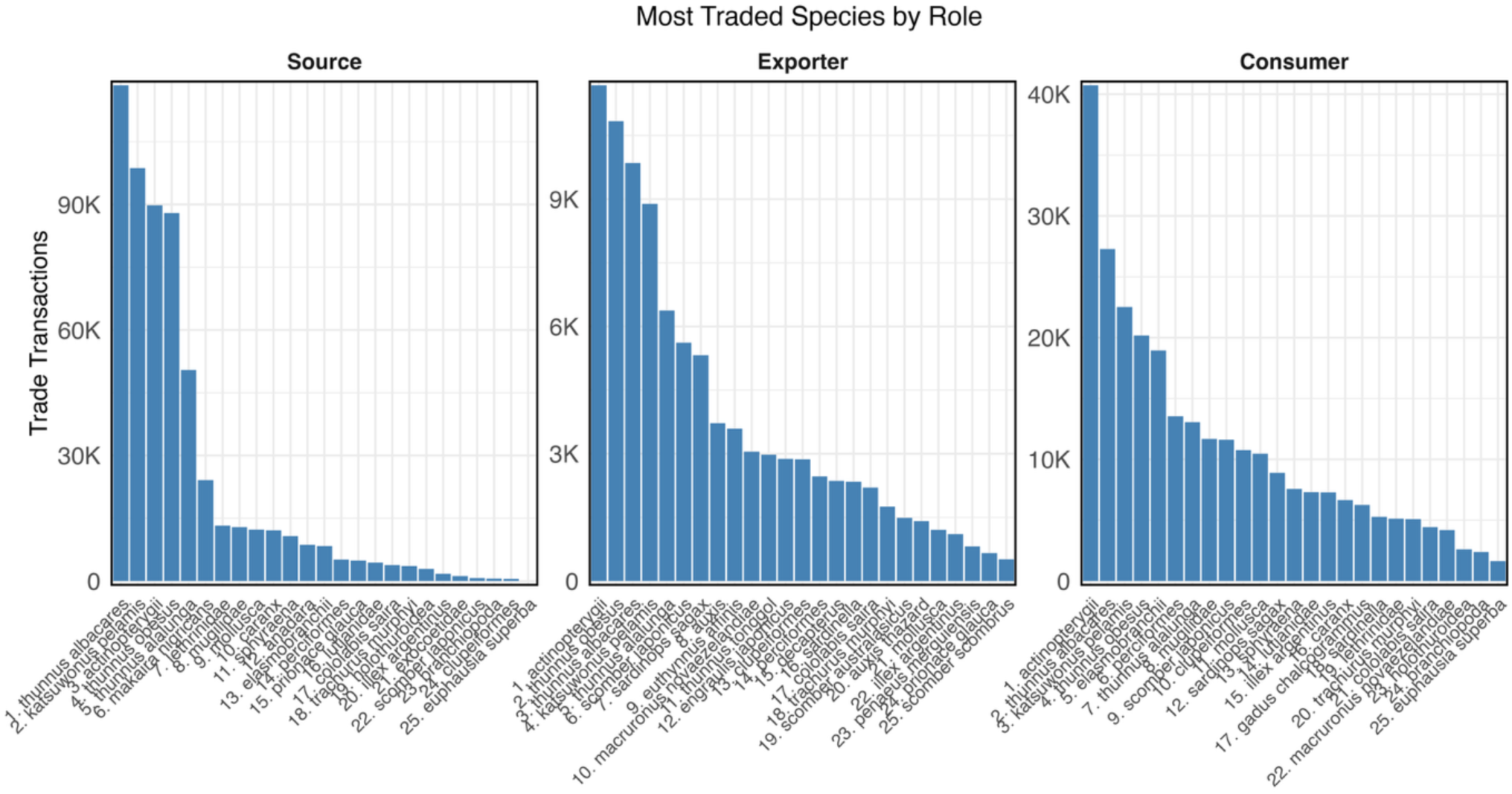
The most traded species for the source, exporter, and consumer country roles in the regional PICs trade network, ranked by total trade transactions.

**Figure S2.**
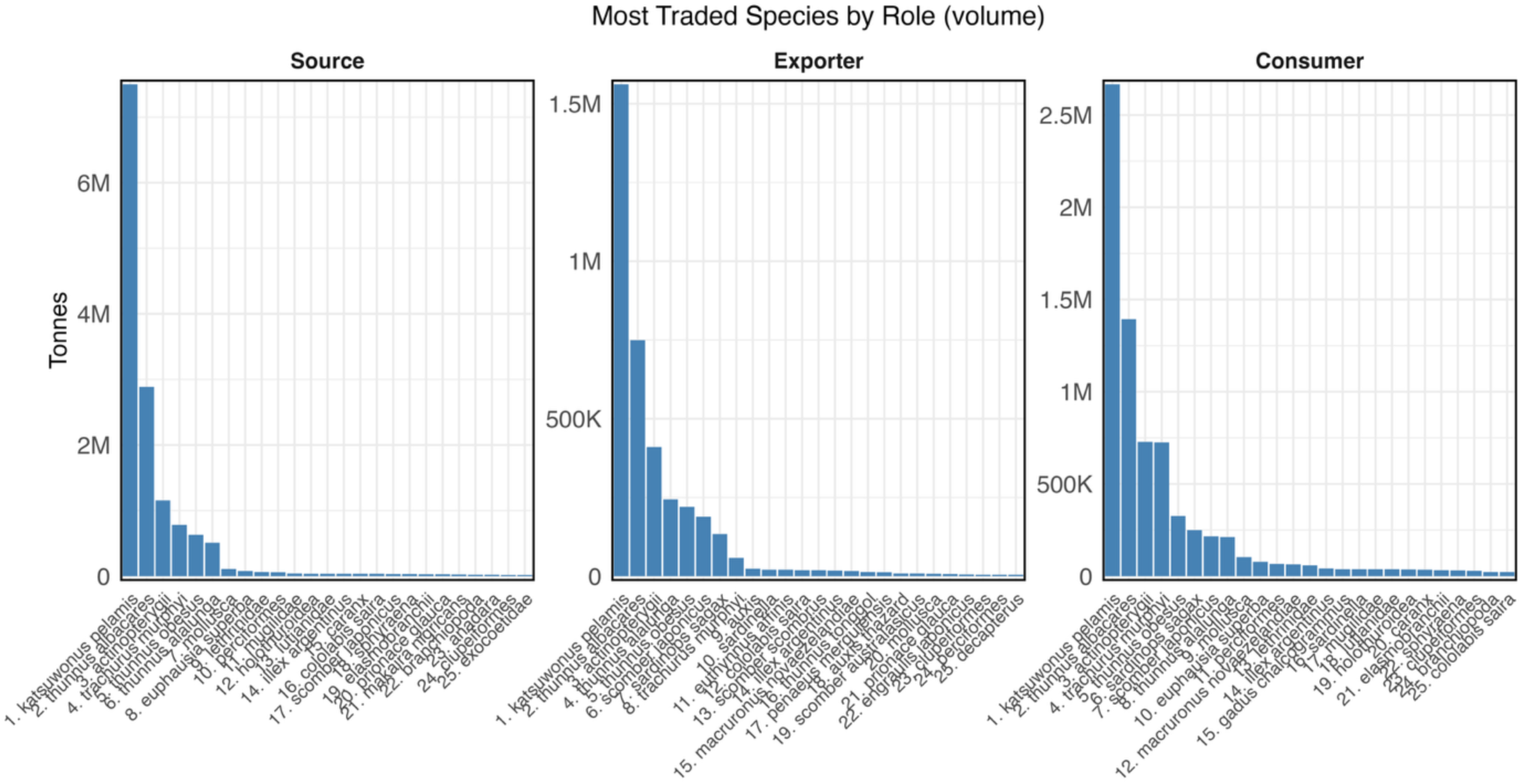
The most traded species for the source, exporter, and consumer country roles in the regional PICs trade network, ranked by total trade tonnage.

**Figure S3.**
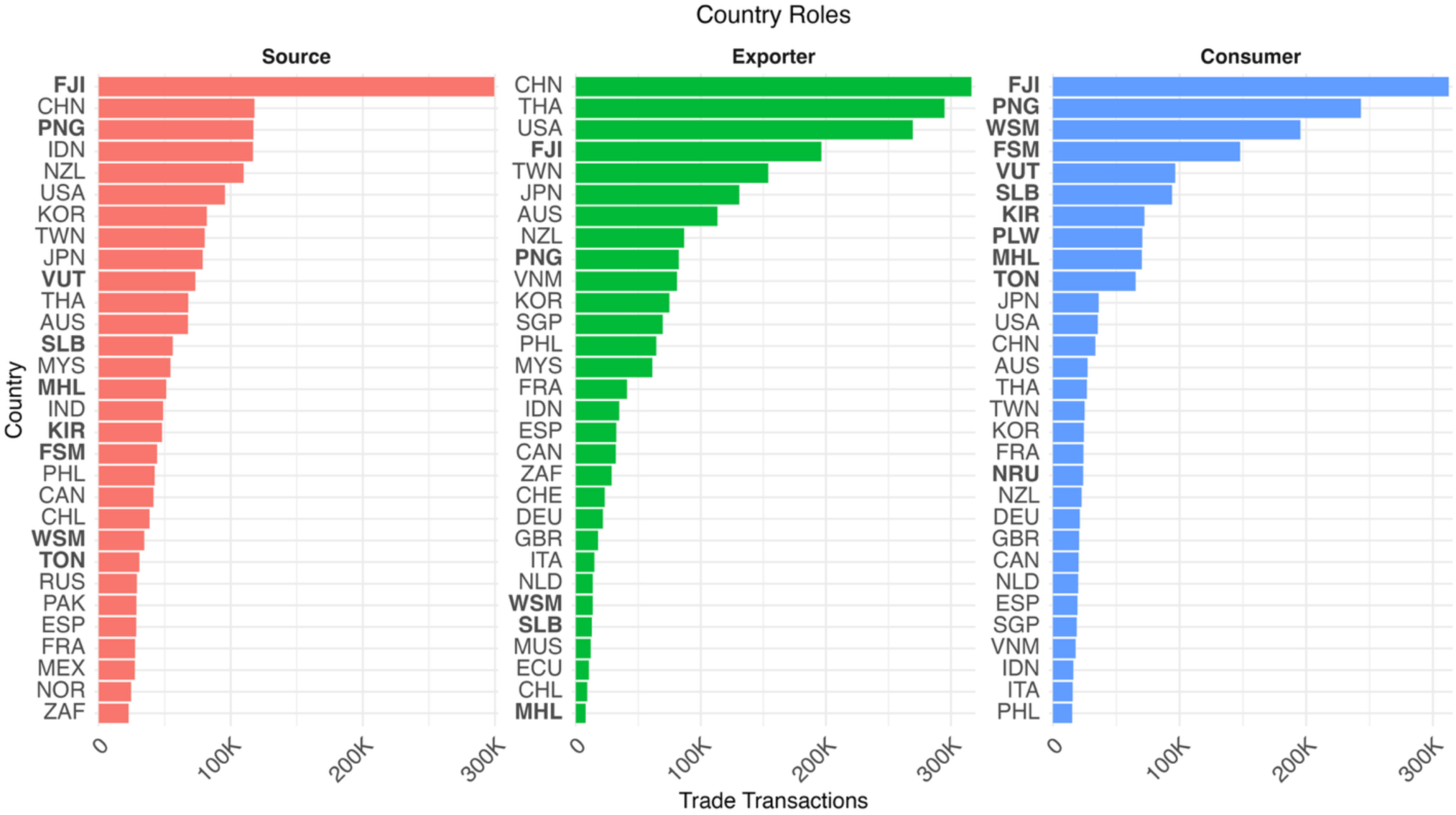
The top 30 countries occupying the source, exporter, and consumer roles in the regional PICs trade network, as measured by number of trade transactions. Countries are indicated with their ISO 3166-1 alpha-3 abbreviations. PICs are bolded.

**Figure S4.**
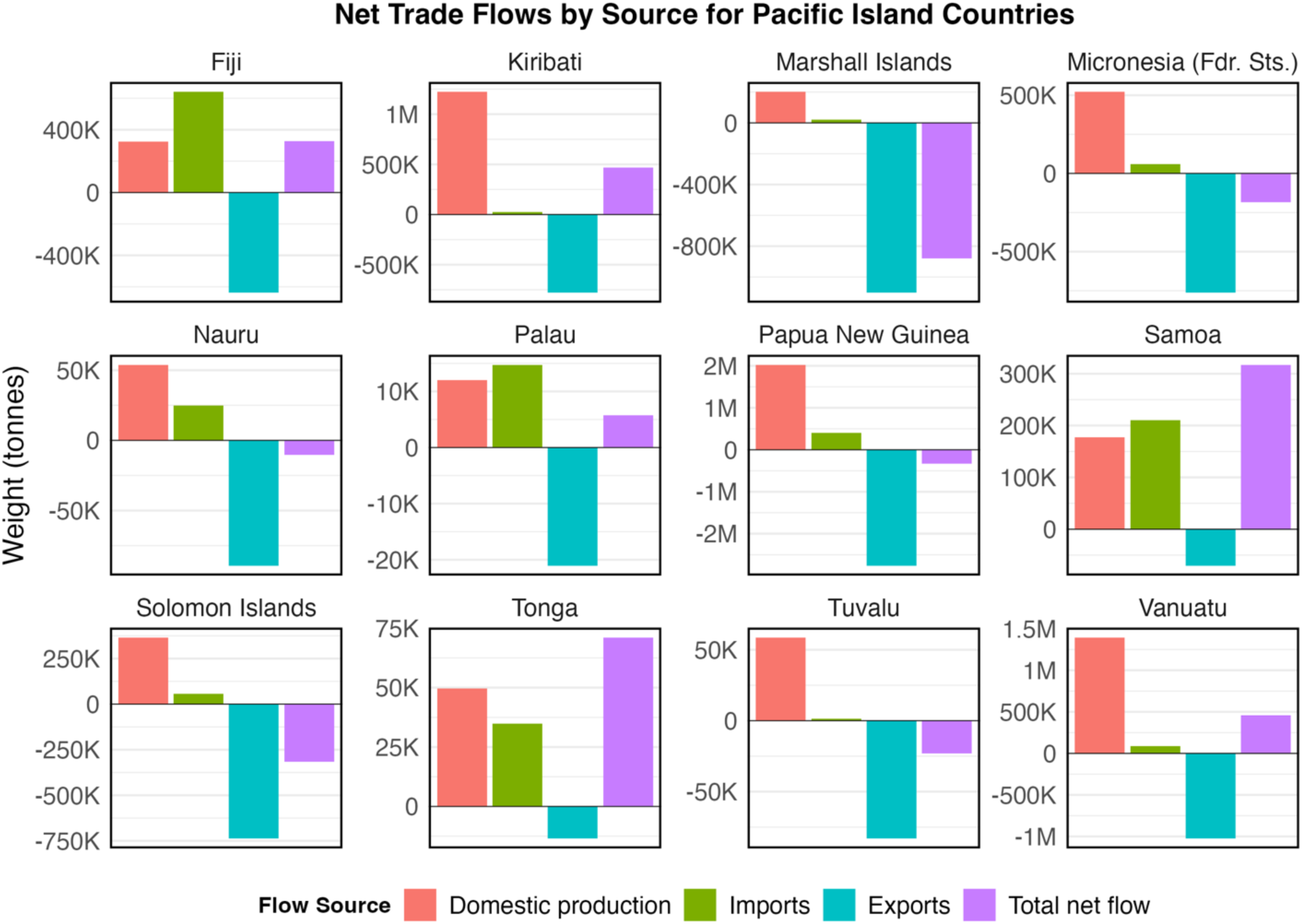
Fisheries trade activity in each PIC, showing how domestic production (red bar), exports (green bar), and imports (blue bar) contribute to overall fish retention or loss. The purple bar reflects the total net flow, which is the sum of the other three flow components. A positive total flow indicates a net surplus of fish retained domestically, while a negative value signals a net loss, meaning more fish were exported than retained in-country.

**Figure S5.**
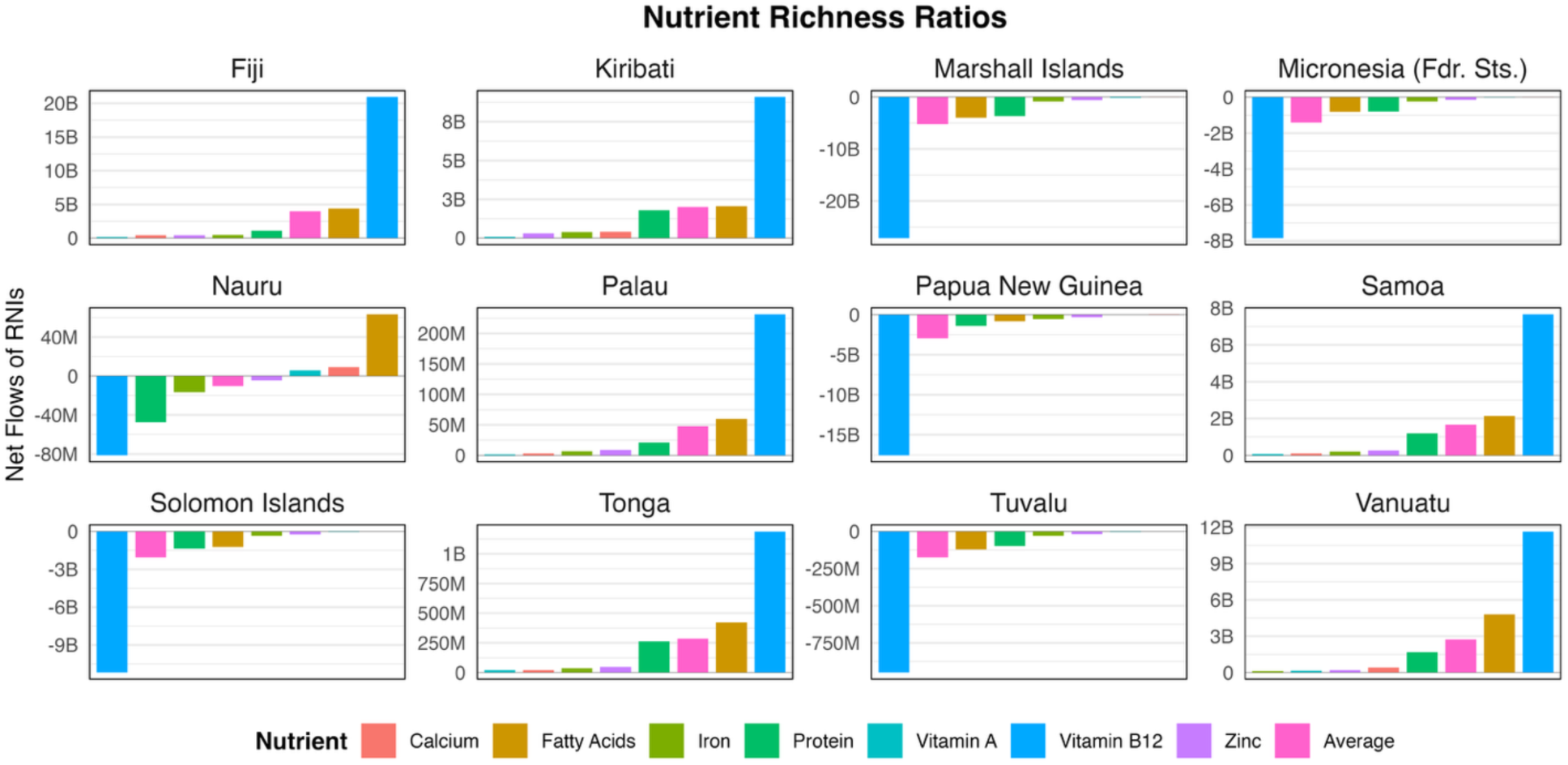
Nutrient richness ratios for each PIC, which represents the net amount of RNIs gained or lost by fisheries trade for each individual nutrient and the average across them. Positive values indicate net retention or in-flow of nutrients, while negative values indicate net losses or out-flow of nutrient

**Figure S6.**
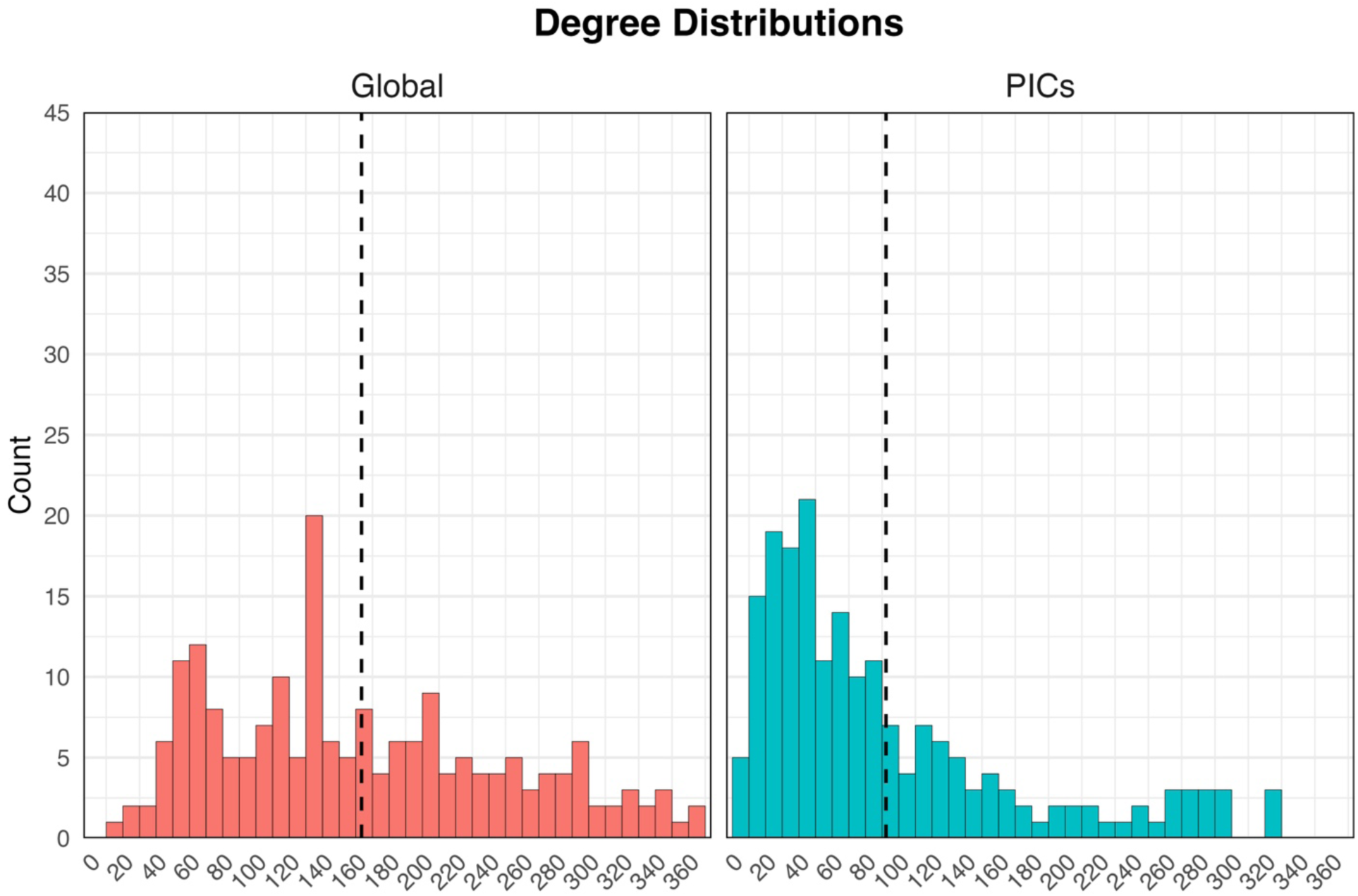
Degree distributions with bin widths of 10 for the global (red) and regional PICs (blue) trade networks. Degrees are the number of edges that a node has. The dashed vertical line indicates the average node degree for each of the network scales (global ∼163, PICs ∼91). Overall, the global degree distribution is somewhat normal, potentially slightly bimodal. The PICs regional network, on the other hand, is slightly more right-skewed, indicating more countries with fewer connections and a generally sparser network.

**Figure S7.**
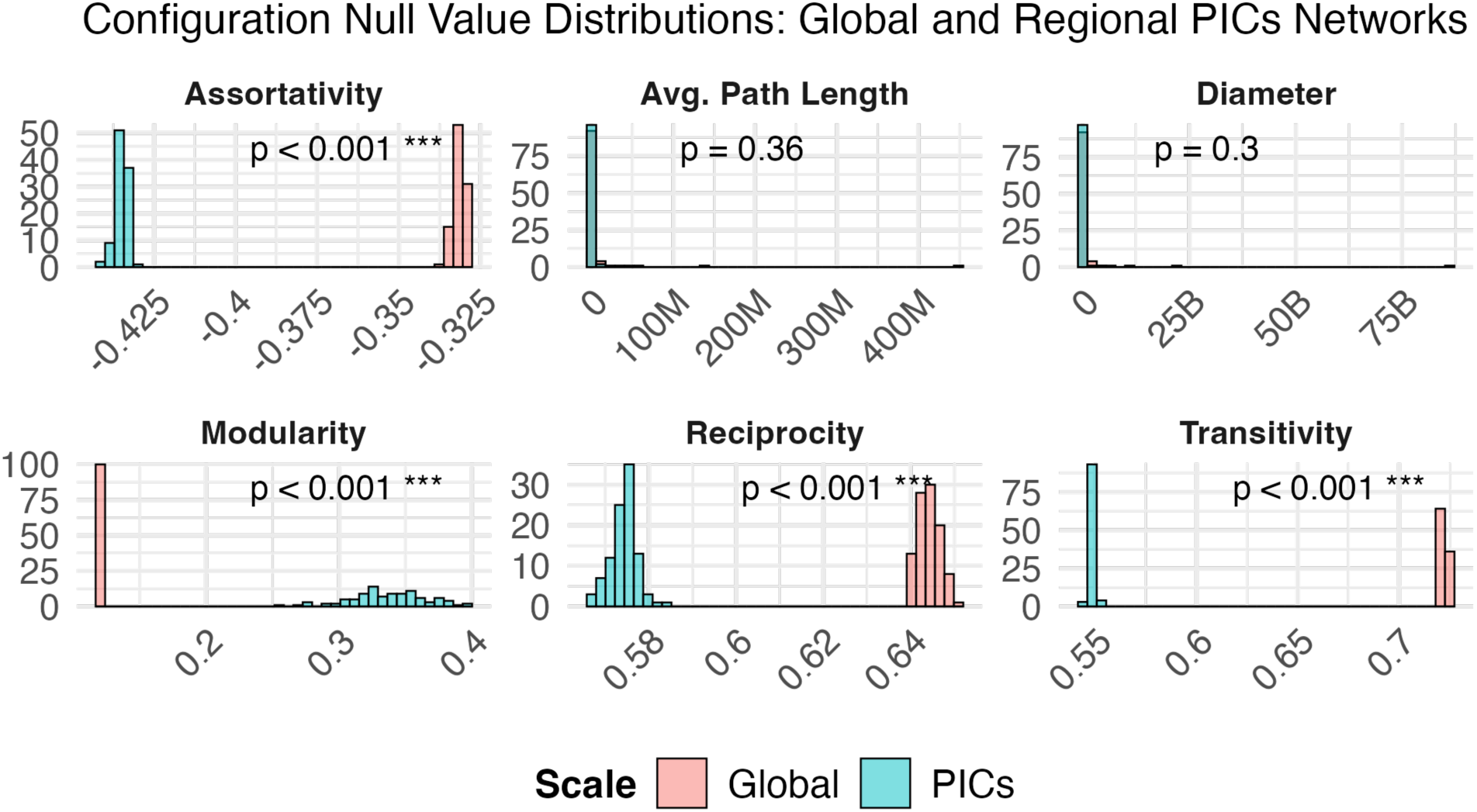
Histogram results for network-wide metrics calculated from the simulated degree-preserving network configuration models built based on the degree distributions of the empirical global and regional PICs nutrient networks. The results for the global network are in pink while the regional results are blue. P-values indicate how statistically different the two models are.

**Figure S8.**
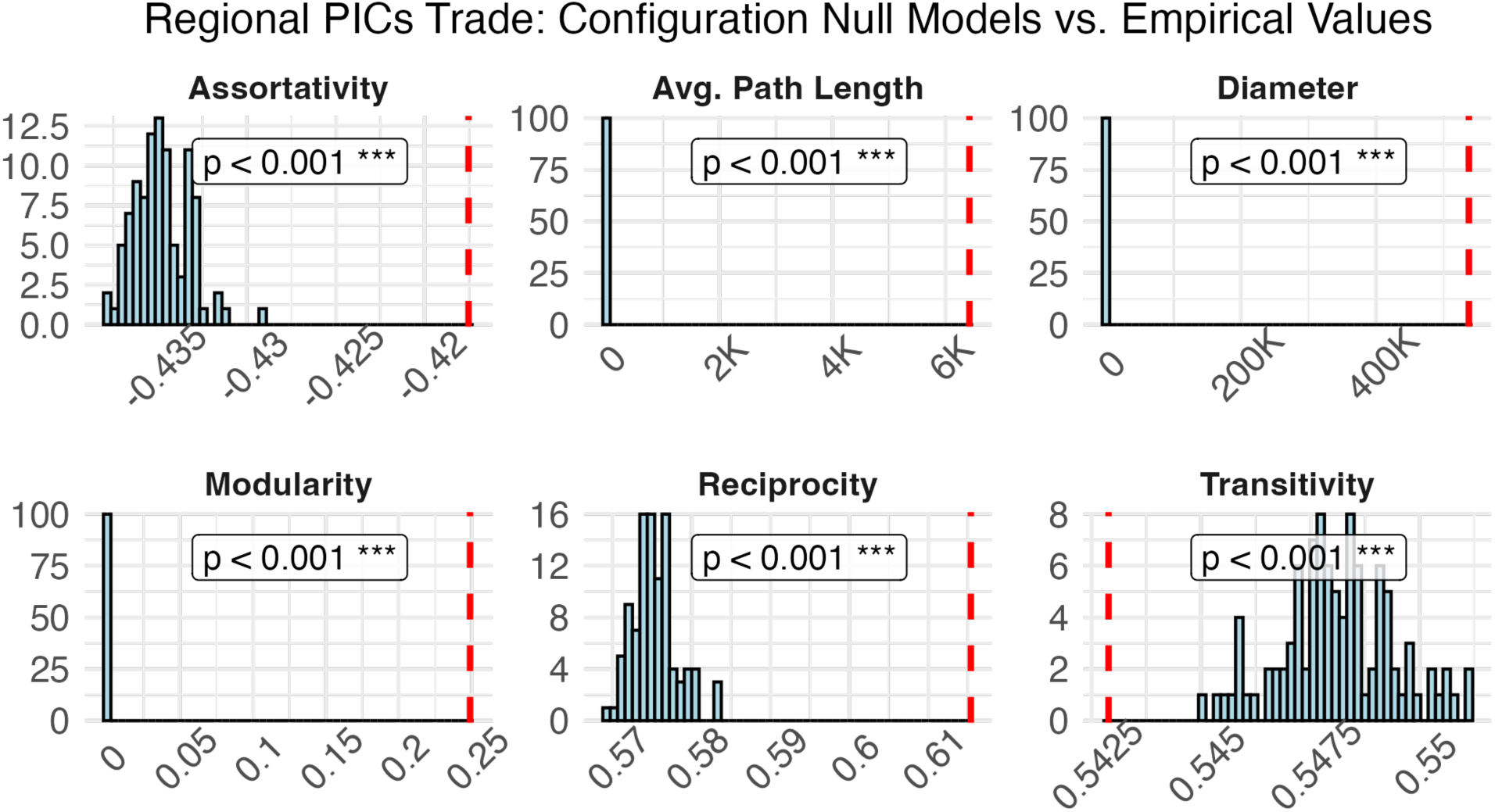
Histogram displaying the simulated network metric values from the repeated degree-preserving configuration model runs (blue histogram) and the empirical network metric values (vertical red dashed line).

**Table S1.** Empirical network metric values for the global and regional (PICs) network.

| **Metric** | **Global** | **Regional** |
| --- | --- | --- |
| Assortativity | -0.29 | -0.42 |
| Average Path Length | 40,641 | 6,422 |
| Centralization | 0.53 | 0.61 |
| Density | 0.43 | 0.24 |
| Diameter | 3,657,939 | 537,208 |
| Modularity | 0.12 | 0.25 |
| Edge number | 15,692 | 8,864 |
| Richness | 192 | 192 |
| Reciprocity | 0.70 | 0.62 |
| Transitivity | 0.71 | 0.54 |

**Table S2.** Two-sample t-test results comparing metrics from the regional PICs and global nutrient flow networks. Average metric values were obtained via repeated network simulations from degree-preserving configuration models based on empirical degree distributions of the regional and global networks. To ensure adequate mixing, we followed convergence guidelines and performed 20×m double-edge swaps before sampling (where m = number of edges), and 2×m swaps between samples. Two-sample t-tests confirmed significant differences across metrics such as assortativity, modularity, reciprocity, and transitivity (all p < 0.001), indicating that regional trade patterns are shaped by factors beyond node connectivity. Centralization and density are fixed by the degree sequence and thus not variable across degree-preserving model runs.

| **Metric** | **Regional Empirical** | **Global Empirical** | **Null Mean (PICs)** | **Null Mean (Global)** | **t-statistic** | **p-value** |
| --- | --- | --- | --- | --- | --- | --- |
| Assortativity | -0.27 | -0.29 | -0.44 | -0.33 | 399.92 | 1.01e-288 |
| Average Path Length | 101,199 | 53,022 | 2,193,957 | 6,545,631 | 0.92 | 0.36 |
| Centralization | 0.089 | 0.49 | 0.61 | 0.53 | - | - |
| Density | 0.16 | 0.45 | 0.24 | 0.43 | - | - |
| Diameter | 3,496,953 | 2,067,148 | 316,282,544 | 1,251,124,907 | 1.04 | 0.30 |
| Modularity | 0.25 | 0.17 | 0.34 | 0.12 | -76.59 | 6.29e-90 |
| Reciprocity | 0.54 | 0.70 | 0.57 | 0.64 | 177.21 | 9.08e-210 |
| Transitivity | 0.55 | 0.74 | 0.55 | 0.72 | 953.68 | 0 |

**Table S3.**
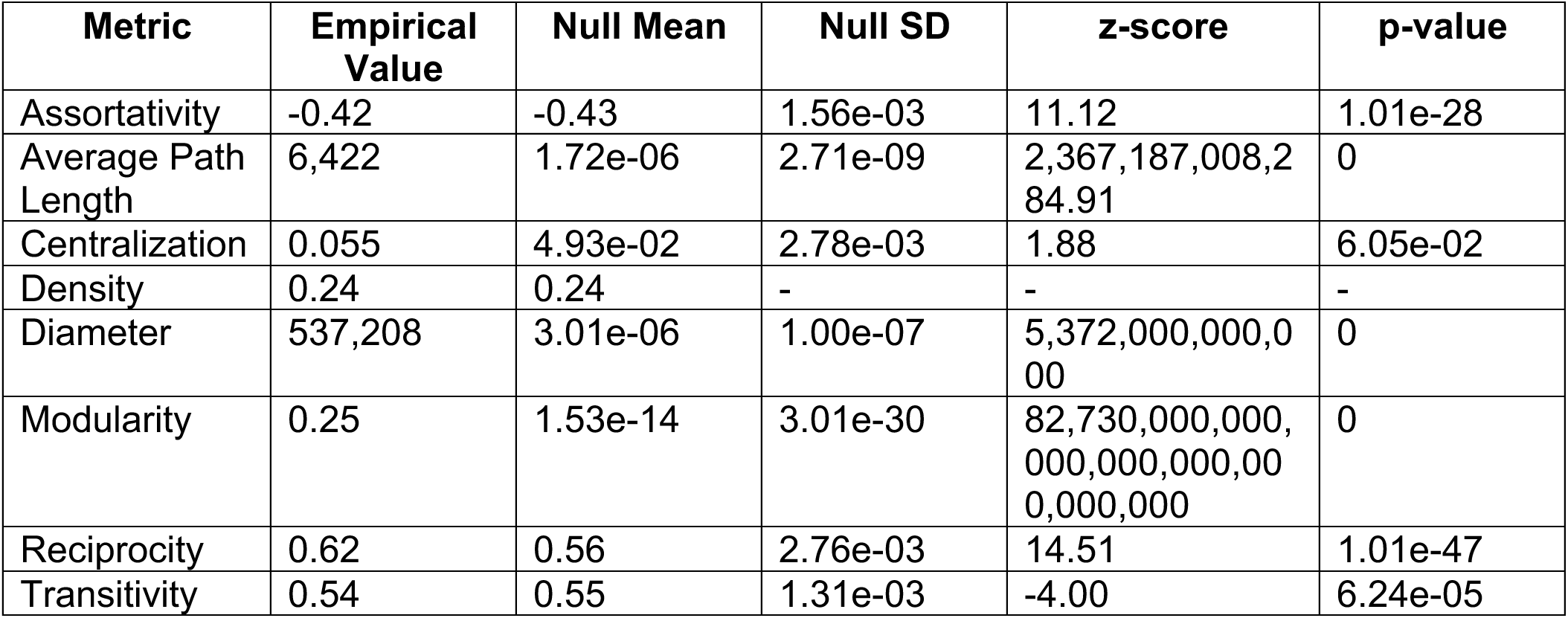
Comparison of empirical regional (PICs) network metrics against 100 null networks generated using a degree-preserving configuration model. Z-scores and p-values reflect how strongly the empirical network deviates from null expectations. Density was not tested due to lack of variation in null simulations. Because the empirical network includes self-loops (e.g., domestic consumption and production), we implemented a custom null model that removed self-loops prior to randomization and reintroduced them post hoc at their original nodes. To ensure adequate mixing, we followed convergence guidelines and performed 20×m double-edge swaps before sampling (where m = number of edges), and 2×m swaps between samples. The PICs empirical network to null models showed several metrics were significantly higher than expected by chance: path length, diameter, modularity, reciprocity, and assortativity (all p < 0.001).

